# Deciphering conformational selectivity in the A_2A_ adenosine G protein-coupled receptor by Free Energy simulations

**DOI:** 10.1101/2021.06.09.447724

**Authors:** Willem Jespers, Laura H. Heitman, Adriaan P. IJzerman, Eddy Sotelo, Gerard J. P. van Westen, Johan Åqvist, Hugo Gutiérrez-de-Terán

**Affiliations:** Department of Cell and Molecular Biology, Uppsala University, Biomedical Center (BMC), Box 596, SE-751 24, Uppsala, Sweden; Drug Discovery and Safety, Leiden Academic Centre for Drug Research, Einsteinweg 55, 2333 CC, Leiden, The Netherlands; Centro Singular de Investigación en Química Biolóxica y Materiais Moleculares (CIQUS); Departamento de Química Orgánica, Facultade de Farmacia. Universidade de Santiago de Compostela, 15782. Santiago de Compostela, Spain; Oncode Institute, 2333 CC Leiden, Leiden; Science for Life Laboratories, BMC, Uppsala (Sweden)

## Abstract

Transmembranal G Protein-Coupled Receptors (GPCRs) transduce extracellular chemical signals to the cell, via conformational change from a resting (inactive) to an active (canonically bound to a G-protein) conformation. Receptor activation is normally modulated by extracellular ligand binding, but mutations in the receptor can also shift this equilibrium by stabilizing different conformational states. In this work, we built structure-energetic relationships of receptor activation based on original thermodynamic cycles that represent the conformational equilibrium of the prototypical A_2A_ adenosine receptor (AR). These cycles were solved with efficient free energy perturbation (FEP) protocols, allowing to distinguish the pharmacological profile of different series of A_2A_AR agonists with different efficacies. The modulatory effects of point mutations on the basal activity of the receptor or on ligand efficacies could also be detected. This methodology can guide GPCR ligand design with tailored pharmacological properties, or allow the identification of mutations that modulate receptor activation with potential clinical implications.

**Author Summary:** The design of new ligands as chemical modulators of G protein-coupled receptors (GPCRs) has benefited considerably during the last years of advances in both the structural and computational biology disciplines. Within the last, area, the use of free energy calculation methods has arisen as a computational tool to predict ligand affinities to explain structure-affinity relationships and guide lead optimization campaigns. However, our comprehension of the structural determinants of ligands with different pharmacological profile is scarce, and knowledge of the chemical modifications associated with an agonistic or antagonistic profile would be extremely valuable. We herein report an original implementation of the thermodynamic cycles associated with free energy perturbation (FEP) simulations, to mimic the conformational equilibrium between active and inactive GPCRs, and establish a framework to describe pharmacological profiles as a function of the ligands selectivity for a given receptor conformation. The advantage of this method resides into its simplicity of use, and the only consideration of active and inactive conformations of the receptor, with no simulation of the transitions between them. This model can accurately predict the pharmacological profile of series of full and partial agonists as opposed to antagonists of the A_2A_ adenosine receptor, and moreover, how certain mutations associated with modulation of basal activity can influence this pharmacological profiles, which enables our understanding of such clinically relevant mutations.

## Introduction

G Protein-Coupled Receptors (GPCRs) are membrane proteins that transduce the signals of hormones, neurotransmitters and metabolites into an appropriate cellular response (1). The canonical intracellular signalling pathways are mediated by heterotrimeric G proteins, though alternative pathways exist like those involving β-arrestin proteins. GPCRs are widely involved in human physiology, where over 800 genes encode six GPCR classes (2) and they constitute the main target of approximately 34% of marketed drugs (3). Our knowledge of the structure-function relationships of GPCRs has increased tremendously in the last decades, largely fuelled by a growing number of GPCR structures. The first crystal structures corresponded to inactive states, following strategies such as fusing the receptor with stabilizing proteins (4) or the introduction of state-specific thermostabilizing mutations (5). The last approach allowed obtaining the first structures in a pseudo-active state in complex with an agonist, where the receptor showed the initial conformational changes characteristic of activation but still lacking any intracellular binding partner(6). The structures of ternary GPCR complexes, including the intracellular signalling G proteins or β-arrestin in addition to an orthosteric agonist, were resolved in recent years mainly due to advances in cryo-EM (7,8). Traditionally, GPCR activation has been described as a two-state model, where the receptor would transition from an inactive (R) to an active (R*) conformation (Figure 1A) (9). This relatively simplistic model becomes more realistic when considering the influence on the receptor equilibrium of chemical modulators (Figure 1B), receptor mutations (Fig 1C) or even the intracellular signalling protein(10).

**Figure 1:**
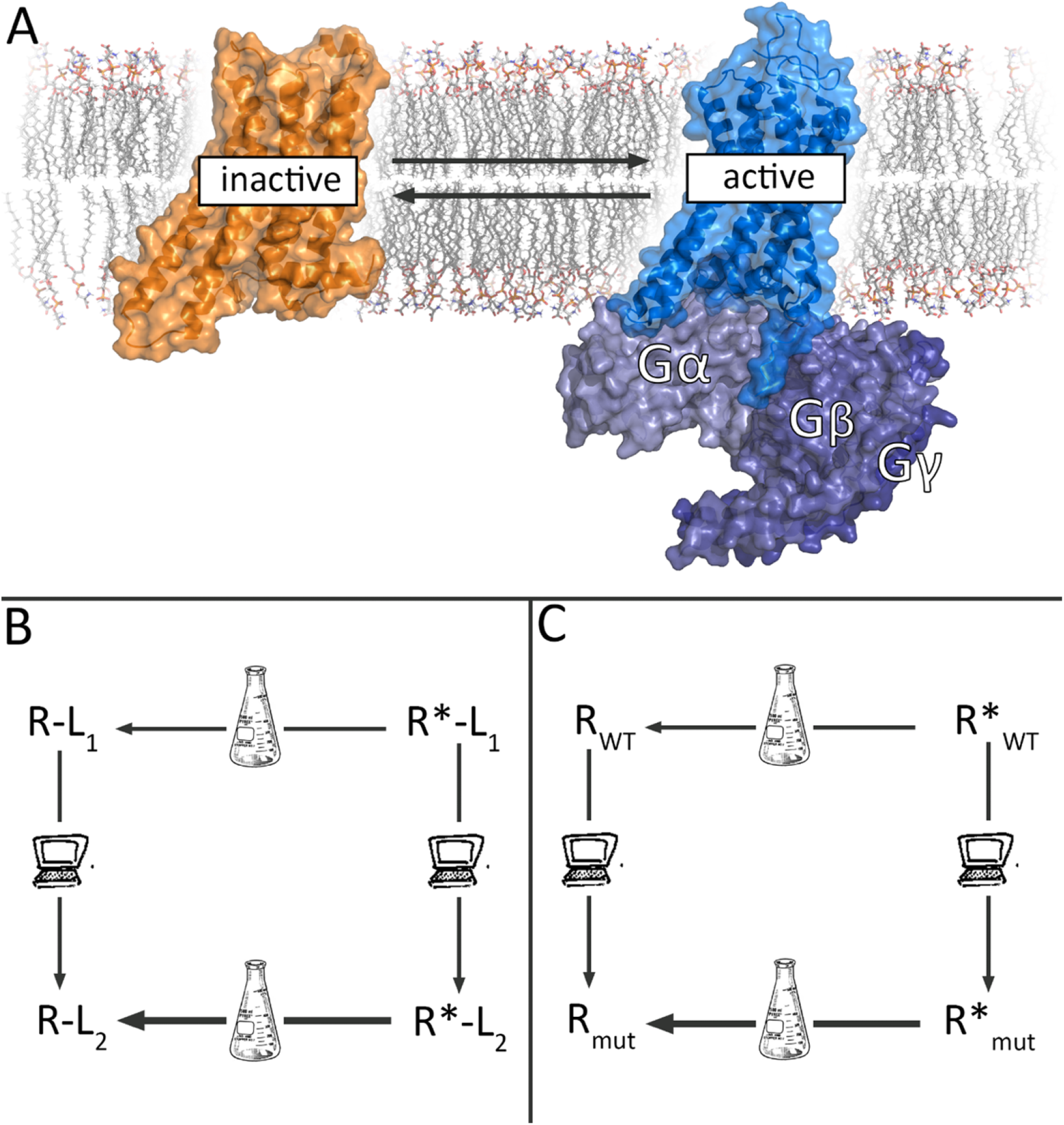
A) Two-state model of GPCR activation between the inactive (orange) and active structure bound to a G protein (blue). B) Thermodynamic cycle linking ligand efficacy to relative affinities for a given receptor conformation. The experimental (horizontal) legs represent the neutral effect of an antagonist (L1) on the receptor equilibrium, and the induced shift towards the active state (R*) due to agonist (L2) binding. C) the receptor basal equilibrium can be shifted by a certain point mutation, in this example a CIM that selectively stabilizes the inactive state. Each thermodynamic cycle can be closed with vertical legs, representing the corresponding FEP calculations between the ligand pair (B) or the receptor mutation (C), allowing to estimate the differences in experimental values as the calculated difference in the FEP legs.

The four adenosine receptors (ARs), namely A_1_, A_2A_, A_2B_ and A_3_, constitute one of the best structurally characterized GPCR families (11). With more than 50 entries in the PDB, the A_2A_AR was one of the first receptors to be captured in the three different conformational states (inactive, active-like, ternary complex) (12–14), and was later accompanied by the inactive and ternary complexes of the A_1_AR (15,16). Structural and mutational data of A_2A_AR revealed that conformational changes associated with activation involve a widening of the ‘ribose pocket’, lined by residues T88^3.36^, S277^7.42^ and H278^7.43^ [upper case numbers refer to Ballesteros-Weinstein numbering (17)] (18). All adenosine receptor full agonists known to date contain a ribose moiety (Figure 2), with the hydroxy substituents forming hydrogen bonds with these residues. As such, the stereospecificity of the ribose group is important in receptor activation. On the other hand, partial agonists are molecules that display a reduced maximum efficacy as compared to the full agonists. In the case of A_2A_AR, partial agonists have been reported bearing as most as one hydroxy substituent, which is supposed to form a single hydrogen bond interaction with one of these residues in the ribose pocket (Figure 2). Even compounds with no hydroxyl can behave as partial agonists (i.e. LUF5833, Figure 2), in which case the mechanistic hypothesis for its partial efficacy could rely on an optimal steric fitting of the phenyl substituent in the enlarged ribose pocket of the active state.

**Figure 2:**
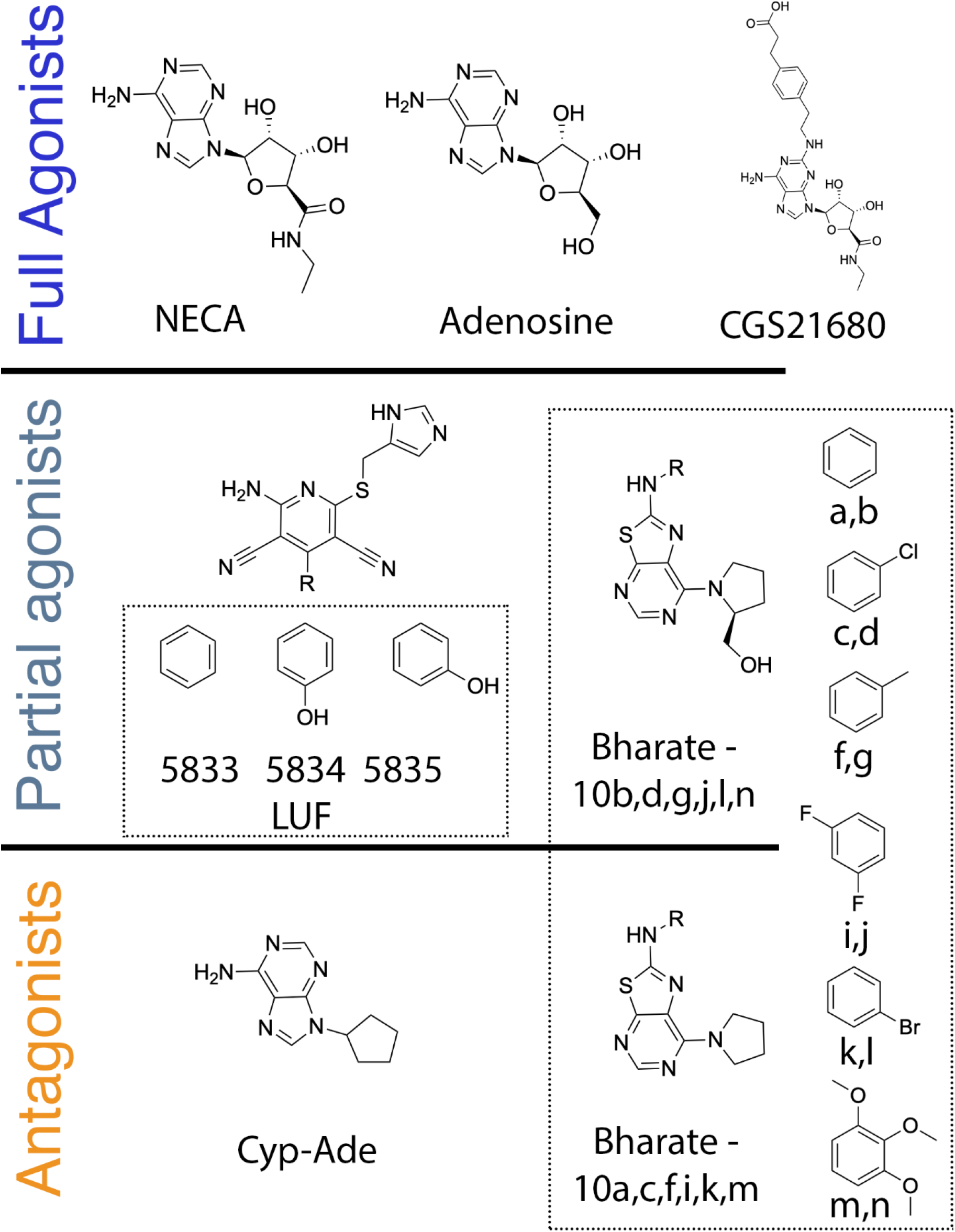
Chemical structures of the compounds considered in this work, classified according to their experimental pharmacological profile.

Site-directed mutagenesis has been traditionally used in GPCR research to characterize the shifts in ligand binding affinities induced by point mutations, and used to assist the elucidation of ligand binding modes as recently reviewed for the AR family (18). Two positions invariably affect both agonist and antagonist ligand binding, namely N253^6.55^ and F168^EL2^, which were later confirmed as anchoring points for all heterocycles present in currently described orthosteric ligands (18). Point mutations can also influence the thermal stability of the receptor, either its overall stability or selectively increasing the stability of one receptor state or conformation (Figure 1C). Moreover, one mutation can simultaneously affect ligand affinity whilst also shifting the receptor conformational equilibrium. For instance, mutations S277^7.42^A and T88^3.36^A, positioned in the ribose pocket, increase the affinity of antagonists while the efficacy of agonists is reduced. In addition to this effect, S277^7.42^A increases the efficacy of some partial agonists (19), while the T88^3.36^A mutation effectively shifts the receptor equilibrium to the inactive state, being classified as a constitutively inactive mutation (CIM) since the basal activity is concomitantly decreased (20). Thus, the overall effect on ligand binding affinity can be a consequence not only of direct interactions with the ligand, but also of the increased stability (and thus availability) of the receptor conformation that preferably binds a pharmacological class of ligand.

Based on the available collection of experimental structures, reliable models of AR-ligand complexes can be generated via computational methods (21,22). While docking algorithms can be very useful for this goal, they typically fail to describe free energies of binding correctly (23). Instead, the increased availability of computational power and algorithms have enabled a routine use of first-principle methods such as free energy perturbation (FEP) to accurately estimate ligand-binding free energies, also for GPCRs (24). In this scenario, we have recently developed robust FEP protocols that were thoroughly applied in the context of GPCR ligand-binding investigations: QligFEP (25) allows to systematically compute relative binding affinity changes between a series of ligands, while QresFEP (26) was designed to evaluate the binding affinity shifts due to single point mutations. The synergistic combination of both approaches recently led to the elucidation of the binding mode of series of A_2A_AR antagonists (27). In this work, we extend the applicability of these methods to study the effects of chemical modifications of ligands, as well as receptor mutations on the conformational equilibrium of the receptor, which we relate to efficiency of the ligand as an agonist modulator. First, we analyse the structure-efficacy determinants of series of full and partial agonists. Thereafter, we elucidate the role of the S277^7.42^A and T88^3.36^A mutations in the associated conformational equilibrium of the A_2A_AR receptor, to finally determine the specific role of these mutations on the efficacy of selected partial and full agonists. The outcome of this study can not only aid the design of chemical modulators with tailored pharmacological properties, but also be broadly applicable to characterize GPCR mutations with clinically relevant effects.

## Results

### Conformational selectivity of ligands depends on their pharmacological profile

In this first section, we explore the predicted conformational selectivity for a number of ligands, as a function of their pharmacological profiles. To do this, a thermodynamic cycle was designed to estimate, for a given molecular pair of e.g. agonist and antagonist, the relative affinities between the two relevant conformational states of the A_2A_AR. The cycle is solved by subtracting the corresponding binding free energies (Δ*G*_b_), calculated via an FEP transformation (agonist → antagonist) performed in the inactive state (Δ*G*_b,R_), from the same transformation performed in the active state (Δ*G*_b,R*_, vertical legs in Figure 1B). In each case, the pair of agonist and antagonist molecules to compare should share enough chemical similarity (e.g. bearing the same chemical scaffold) while maintaining a sufficiently distinct pharmacological profile. This scheme was consequently used to investigate the variability in the pharmacological profile within three chemical scaffolds: *i)* the classical ribose-containing agonists, such as NECA; *ii)* 2-phenylaminothiazolo[5,4-d]pyrimidines, which were recently characterized as partial agonists depending on the substituent in position 7 (28); and *iii)* partial agonists derived from the 4-phenylpyridine scaffold of LUF5834 (29). In each case, the (partial/full) agonist was compared to a chemical analog that behaves as a (neutral) antagonist (Figure 2).

The experimentally determined coordinates of classical agonists binding to the A_2A_AR (Figure 3A and 3B) were used as a starting configuration for the active-state simulations (denoted with an asterisk), while the corresponding binding mode to the inactive A_2A_AR was generated by receptor superposition. The binding mode of 9-cyclopentyadenine (Cyp-Ade) antagonist where the ribose group of adenosine is replaced by a cyclopentane(30), was generated using a flexible ligand alignment strategy (see Methods). The initial configuration of the two partial agonist chemotypes was generated via docking to the A_2A_AR*, showing occupancy of the ribose pocket (Figure 3C and 3D), while the corresponding antagonist analogues were modeled in an analogous binding orientation in the A_2A_AR inactive structure. All compounds shared similar interactions with N253^6.55^ and F168^EL2^, irrespective of the receptor conformation or ligand chemotype.

**Figure 3:**
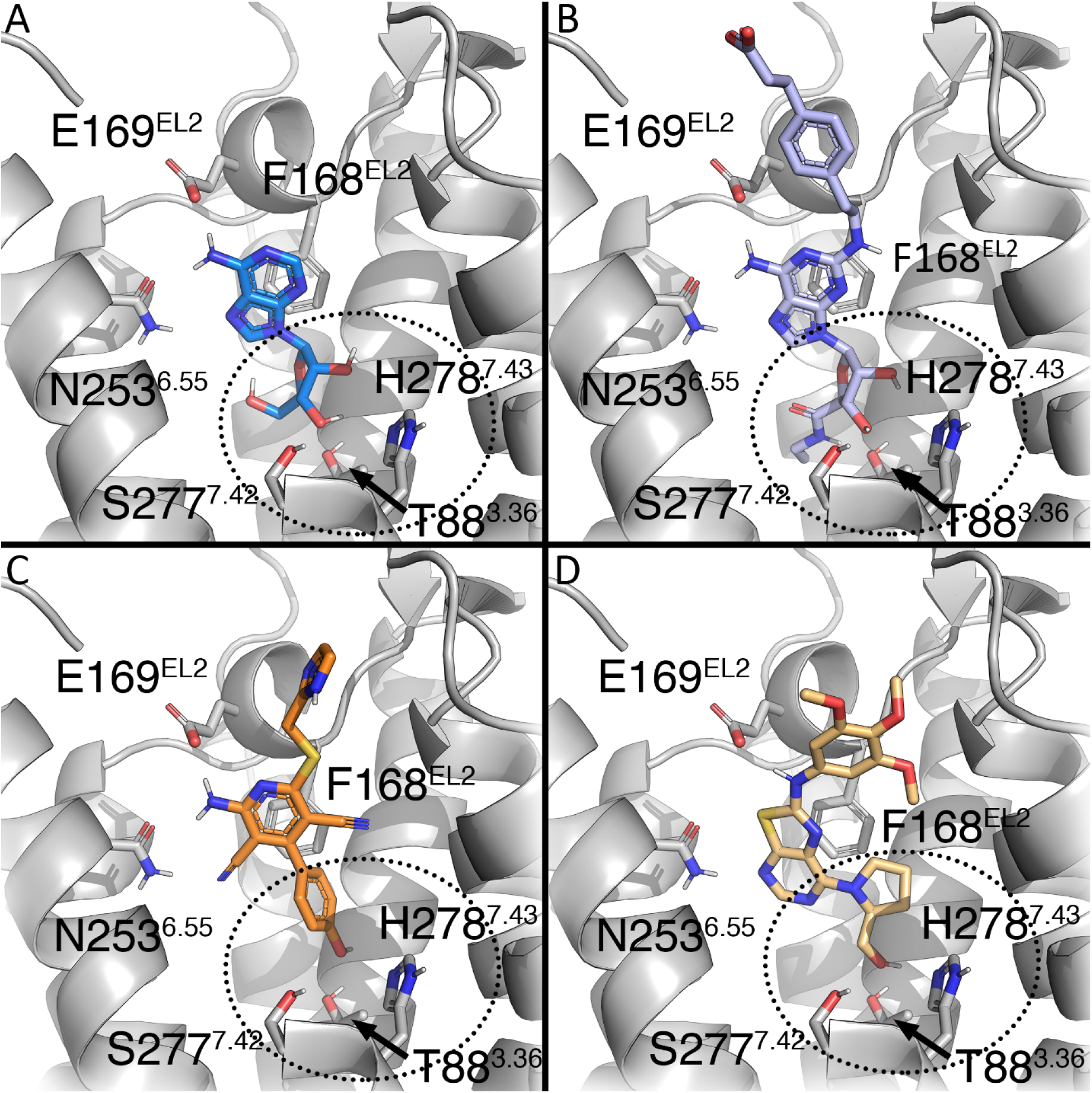
Binding mode of four adenosine A_2A_AR agonists with different efficacies. The top panels show the experimental pose of full agonists **adenosine**(A, PDB code 2YDV (31)) and **CGS21680**(B, PDB code 4UG2(13)). The bottom panels show the docking model obtained for partial agonists **LUF5834**(C) (29) and **10n**(D) (28). The key residues common for ligand-receptor (N253^6.55^, E169^EL2^ and F168^EL2^) (18) are shown in sticks. The ribose binding site is denoted in a dotted circle, with interacting residues associated with ligand activation shown in sticks.

The full agonists NECA and adenosine (ADO) show different experimental efficacies for the activation of A_2A_AR. Consequently, NECA was set as a reference with a maximum efficacy of 100%, with adenosine having 45% efficacy relative to NECA(32), and the neutral antagonist Cyp-Ade having 0% efficacy. A qualitative descriptor, Δ-efficacy, can be defined as the positive difference in % efficacy values between pairs of (partial) agonist and antagonist compounds. In analogy, the thermodynamic cycle depicted in Figure 1B allows to estimate the relative preference for the active conformation for the same pair of molecules, a property that we will try to correlate with the corresponding efficacy shifts. Here, one has to note that, due to their fundamentally different formulation, a full quantitative correlation between the experimental (Δ-efficacy, percentage) and the calculated (ΔΔ*G*, logarithmic) descriptors is not expected. However, a qualitative correlation would indicate that our end-state modeling of ligand efficacy would be useful to explain efficacy shifts between molecule pairs, and thus be of potential interest to further predict ligand pharmacological profiles. In other words, the differences in the calculated ΔΔ*G* might be used to classify the compounds on the basis of their expected efficacy gain from a reference molecule. According to the scheme in Figure 1B, an expected positive value in the calculated ΔΔ*G* would indicate the expected preference of the agonist for the active state. This was indeed the case for the NECA and ADO perturbations to the antagonist Cyp-Ade (Figure 4). Moreover, the magnitude of the calculated ΔΔ*G* value is double for the NECA transformation as compared to the case of adenosine, which indicates the correct ranking of this class of compounds according to their expected efficacy.

**Figure 4:**
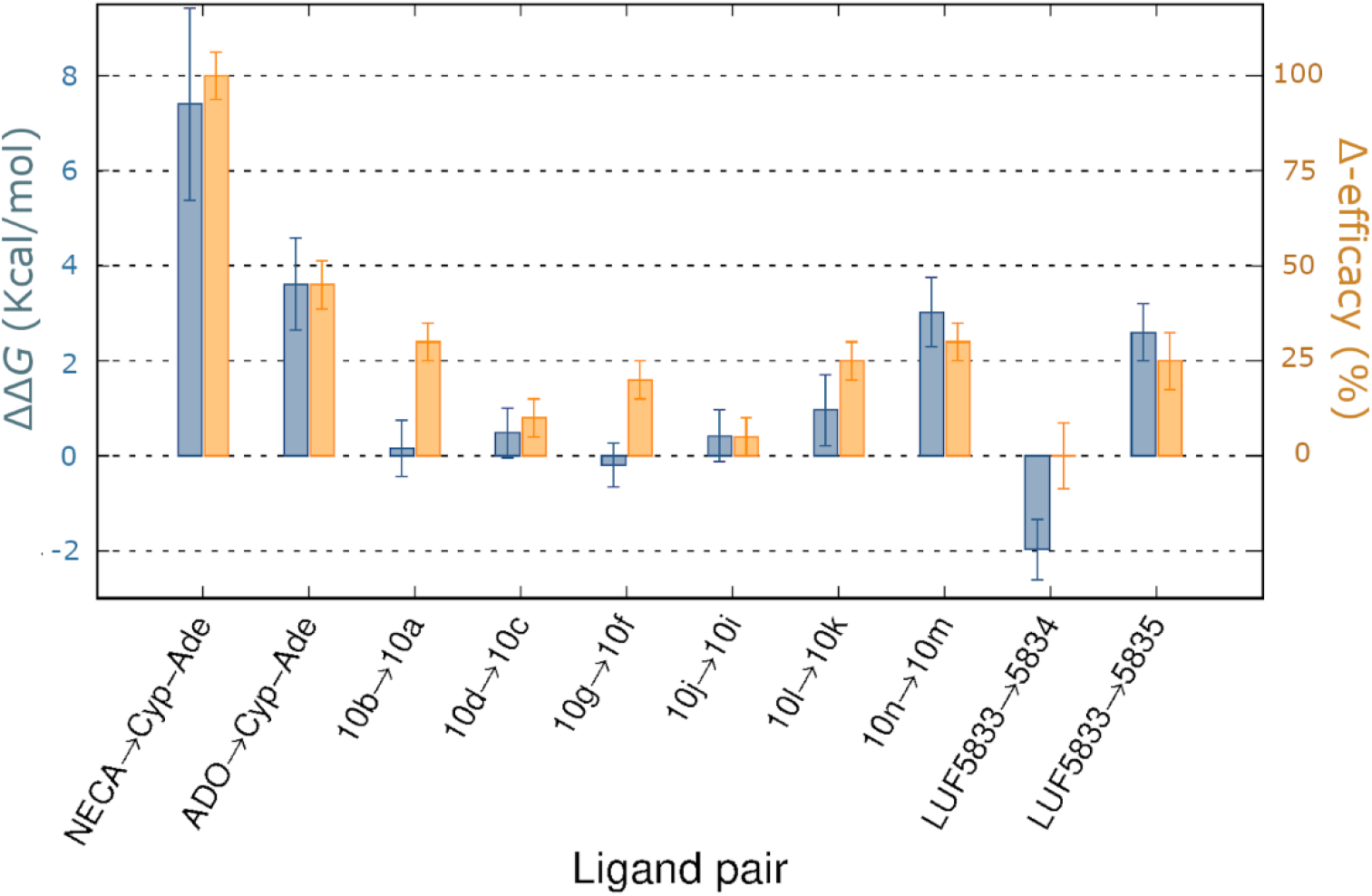
Calculated relative affinities for the active *vs* inactive receptor conformation (ΔΔ*G* = Δ*G*_b,R_ - Δ*G*_b,R*_, kcal/mol, blue bars) for pairs of agonists / antagonist of analogous chemotype (blue bars, average values ± SEM). For each ligand pair, the relative shift in experimental efficacy is shown as Δ-efficacy (orange bars, E_max,ago_-E_max,antago_, in % ± SEM), with corresponding Emax values reported relative to the reference full agonists NECA(28), or CGS21680-for the LUF chemotype (29), see text.

We then moved on to examine the molecular determinants of the agonistic properties for a series of 7‑(prolinol‑N‑yl)-2-phenylaminothiazolo[5,4-d]pyrimidines on the A_2A_AR (28). The SAR and earlier molecular modeling of these series revealed that the prolinol substituent was essential to maintain the partial agonist profile, presumably by partially mimicking the ribose interactions (see compounds, **10b, 10d, 10g, 10j, 10l, 10n,** Figure 2), in contrast to the corresponding pyrrolidine substituted compounds (**10a, 10c, 10f, 10i, 10k and 10m**, Figure 2) which are all neutral antagonists (i.e. with efficacy not significantly different from 0%) (28). Following our preliminary results on this chemotype (33), we systematically applied the thermodynamic cycle in Figure 1B to all corresponding partial agonist/antagonist pairs within this series. A deeper look into the experimental data shows that the efficacy of the partial agonists is modulated by decorations of the exocyclic amino group (R, see Figure 2). Our calculations indicate that compound **10m** would show the highest preference for the active state of the receptor, followed by compound **10k**(Figure 4), in line with their efficacy rank experimentally observed. Noteworthy, in all cases the prolinol-containing compounds displayed a distinct conformational selectivity for the active receptor, as compared to their pyrrolidine substituted compounds, explaining the role of this substituent in the partial agonist profile of this chemotype (Figure 4). The structural interpretation of this data reinforces the role of the hydroxyl group in the 7-prolinol in engaging the residues that trigger A_2A_AR activation (Figure 3).

The third chemotype here explored is derived from the 4-phenylpyridine scaffold, characteristic of a family of A_2A_AR partial agonists represented by LUF5833 (Figure 2) (29). Hydroxy decorations on the phenyl ring have been shown to affect ligand efficacy: the *p*-OH-phenyl in LUF5834 is equipotent to the unsubstituted LUF5833 (55% ± 15% as compared to the efficacy of the reference full agonist CGS21680), while the *m*-OH-phenyl in LUF3835 results in an increased value of this relative efficacy of 80% ± 10% (see Figure 2) (19). In their original report, Lane et al. postulated that these compounds bind with the phenyl substituent deep in the binding pocket of the A_2A_AR, making different interactions with activation-related residues as compared to ribosidic agonists (19). We herein examined if such a binding mode hypothesis (Figure 3) could explain the differences in ligand efficacy following the FEP approach outlined in Figure 1B. In this case, the dehydroxylated partial agonist LUF5833 was used as the reference ligand in two pair comparisons. Since the three compounds show comparable experimental binding *affinities* for the A_2A_AR, the experimental increase in *efficacy* for a given derivative can be directly correlated with its increased relative affinity for the active (R*) over inactive (R) receptor conformation, which is precisely the output of our calculations. The results (Figure 4) indicate that the introduction of a *m*-OH-phenyl in LUF5835 would lead to a significant conformational selection for the active receptor, in line with the experimental gain of approximately 30% efficacy(19). The structural interpretation of this predicted efficacy is located on the hydrogen bond between the OH group in *meta* position of LUF5835 with T88^3.36^, observed in the simulations with R*, an interaction that is well known to be involved in agonist recognition(18). However, the introduction of a *p*-OH-phenyl substitution in the reference ligand leads to a reduced predicted efficacy, since this substituent would not be making any preferred interaction in the active state as compared to the inactive, with the experimental data showing equipotency of this compound pair.

During the preparation of this manuscript, a new crystal structure of compound LUF5833 in complex with the inactive conformation of the A_2A_AR was published (34). Comparison of our model with this structure revealed an almost identical inactive conformation for the receptor (RMSD_A2AAR_ = 0.78 Å). The ligand position was also in good agreement with our MD simulations with an overall RMSD_LUF5833_ = 2.20 ± 0.45 Å, calculated over the last 10% of the FEP trajectories. Specifically, most variability was located on the flexible 1H-imidazol-2-ylmethylsulfanyl substituent oriented towards the extracellular cavity, with the core moiety being more stable along the MD simulation (RMSD_LUF5833_core_ = 1.38 ± 0.51 Å). The structure also demonstrates that the partial agonist LUF5833 can actually bind to the inactive conformation of the receptor, which supports the line of reasoning of this study.

### The effect of point mutations on basal activity and conformational selectivity

According to the experimental mutagenesis data, the activation trigger induced by the partial agonist LUF5834 would involve different interactions with residues in the ribose binding site, as compared to ribose-containing full agonists (19). In that study, CGS21680, a C2-substituted variation of NECA (Figures 2 and 3), was used as a reference full agonist in the pharmacological characterization of the LUF series of compounds. Particularly intriguing was the effect of two mutations, T88^3.36^A and S277^7.42^A, on the modulation of the efficacy of these two molecules. While the binding affinity and potency of CGS21680 was severely reduced by both mutations, the potency of LUF5834 was unaltered or even slightly increased (19).

Consequently, we wondered about the molecular mechanism of the different mutational-induced shifts on ligand efficacies. This is indeed a complex question, as one can imagine two mechanisms via which a mutation can modulate the efficacy of an agonist: On the one hand, the mutation might affect the basal activity of the receptor, by selectively stabilizing one conformation. In this case, while the T88^3.36^A mutant is a CIM that reduces the basal activity (19), by selective thermal stabilization of the antagonist-bound conformation(20), no significant effect was observed on the basal activity for the S277^7.42^A mutant (19). On the other hand, the same mutation can directly modulate the binding affinity of the (partial) agonist for the effective, active conformation. While there is no experimental data for the conformational affinity, our approach allows to model each of these effects independently and combine them a posteriori, taking advantage of the possibility of combining two thermodynamic cycles if they share a common leg.

As shown in Figure 1C, the effect of a mutation on the predicted basal activity can be modeled by designing a thermodynamic cycle that represents the effect of the corresponding Ala mutation on the conformational selectivity. The cycle can be solved through FEP simulations of the vertical legs, i.e. annihilation of the sidechain of interest (wt → Ala perturbation) in each receptor state (R, inactive and R*, active). Such model accounting for mutational effects on GPCR *basal activity* can be combined with a second thermodynamic cycle, accounting for the mutational shifts in ligand binding *affinity* for the *active* conformation, which we extensively used to explain mutagenesis effects on A_2A_AR agonist binding (35). The addition of these two cycles (Figure 5A) should yield as a net result the effect the mutation on the ligand internal *efficacy*, estimated as the difference between the two vertical legs delimiting the combined thermodynamic cycle in Figure 5A, i.e. 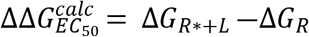 (Figure 5B, solid blue columns). It should be noted that, in this case, the experimental parameter that we are trying to match with the relative free energy calculations is not the maximum efficacy (%E_max_), but instead variations in the internal efficacy of each compound (EC_50_). Thus, a numerical (quantitative) correlation between calculated and experimental values can in this case be attempted, by expressing the experimental shift in EC_50_ values induced by a mutation as 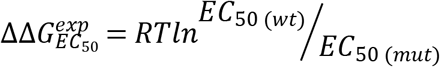 (Figure 5B, solid orange columns). Finally, the additional calculation of the shared vertical leg, Δ*G*_R*_, which is canceled in the combined cycle, allows to separate the effect on basal activity (ΔΔ*G*_R*_, Figure 5A, left side, Figure 5B, dense dashed columns) and on ligand affinity for R* (ΔΔ*G*_L-R*_, Figure 5A, right side; Figure 5B, light dashed columns). As we will see, this can provide valuable information for the interpretation of the calculated data.

**Figure 5:**
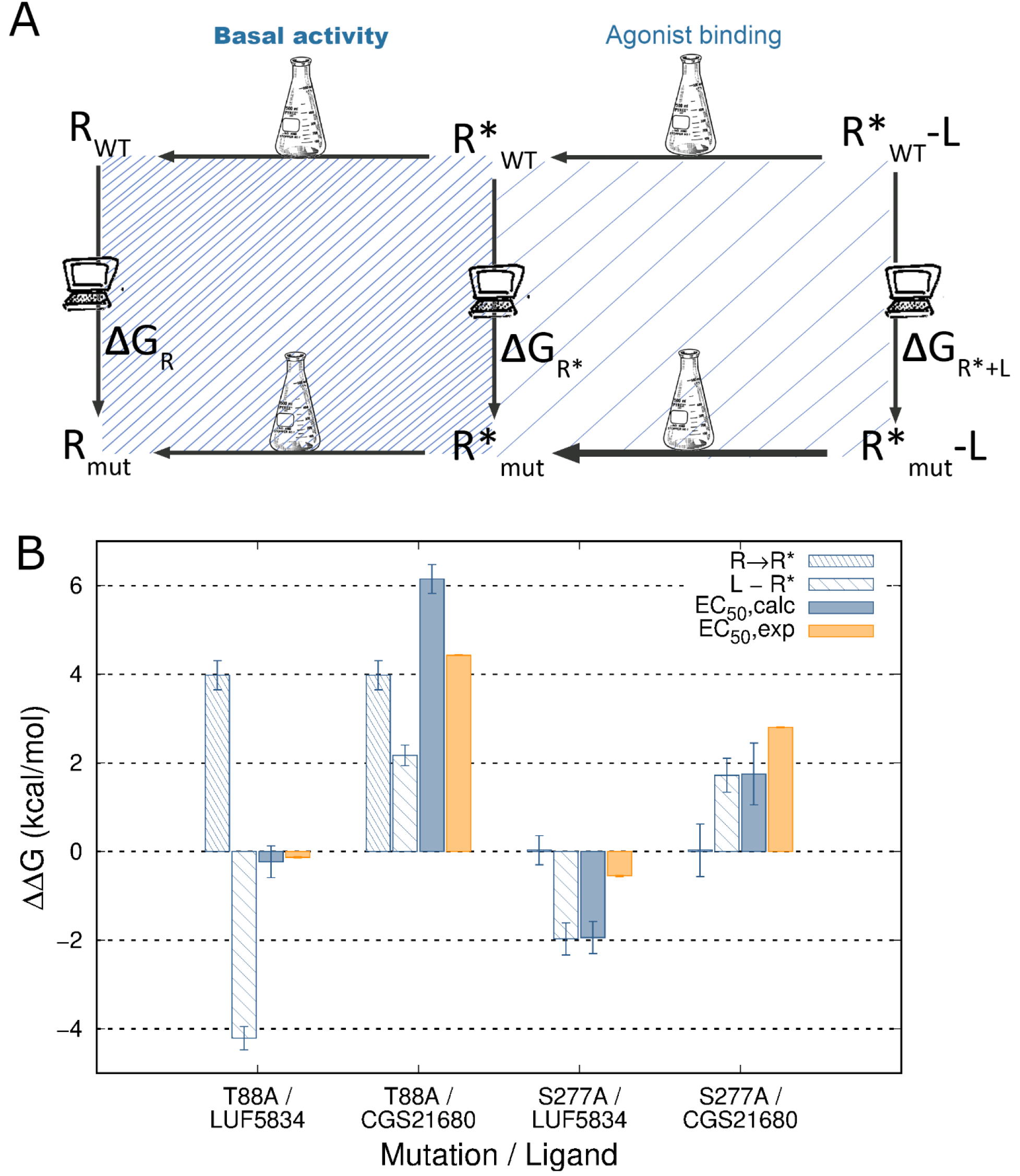
(A) Thermodynamic cycles representing the effect of a point mutation (mut) on the distribution of inactive (R) and active (R*) states of the receptor (left side, dense dashed), and on the affinity of a ligand (L) for the active state (right side, light dashed). The combination of the two thermodynamic cycles would yield the net effect of the mutation ligand efficacy. (B) Calculated effects of point mutations on the A_2A_AR constitutive activity (ΔΔG_R* →R_, dense dashed columns), and on the ligand relative affinity for R* (ΔΔG_L-R*_, light dashed columns), following the corresponding thermodynamic cycles depicted in (A). The overall effect of the mutation on the shift in ligand efficacy, (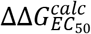, solid blue) is calculated by combination of these two values, showing correlation with the experimental values (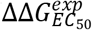, solid orange, see text).

We first looked into the effects on the basal activity of the receptor of the T88^3.36^A and S277^7.42^A mutations. According to our model, the CIM, T88^3.36^A should increase the relative stability of the inactive receptor, which is precisely the outcome of the corresponding FEP simulations (Figure 5B). The active state of the T88^3.36^A mutant is significantly less stable than the wt, with ΔΔ*G*_R* →R_ = 3.98 kcal/mol (Supplementary Table 1). In contrast, a similar analysis of the S277^7.42^A mutation shows a negligible value for the calculated values of ΔΔG_R* →R_, (Figure 5B, Supplementary Table 1), meaning that the conformational equilibrium should not be affected by the mutation, in line with the experimentally observed neutral effect of this mutation on the basal activity of the receptor (19).

The overall mutational effects on the modulation of ligand efficacies were then calculated and compared to the experimental efficacy shifts in each case (Figure 5B, solid columns). One can observe that the modeled T88^3.36^A mutation does not significantly affect the predicted internal efficacy of LUF5834, in line with the experimental data (19). A deeper look at the data allows us to envisage a mechanism for this neutral effect, since the relative increase in affinity for the active state is compensated by a decrease in the population of this active state induced by the mutation (thin and thick dashed bars, respectively, in Fig 5B). Conversely, the model indicates that the S277^7.42^A mutation does not affect the conformational equilibrium of the receptor, and in this case the predicted increase in affinity of LUF5834 for the active state is translated into a net increase of the efficacy of this ligand upon mutation, in agreement with the experiment data (19). For the full agonist CGS21680, both mutations resulted in a similar decrease in estimated ligand affinities, in agreement with our previous computational modeling of the mutagenesis data for ribose-containing ligands(35). However, this effect is amplified by the diminished availability of the active state upon T88^3.36^A mutation, resulting in a much drastic estimated reduction of CGS21680 efficacy as compared to the S277^7.42^A mutant. These results allow a correct ranking of this pair of mutations in terms of how they affect the efficacy of this ligand, in line with the experimental observation where the reduction in the efficacy of CGS21680 is 10 times higher for the T88^3.36^A mutant (19).

## Discussion

The use of FEP simulations coupled to a combination of thermodynamic cycles to study the effect of mutations on binding and catalysis of subtilisin was first presented in a seminal paper of Rao *et al.* in the late eighties(36). Starting from that idea, in this work we introduce a novel approach to estimate the modulation of GPCR activation, based on an original design of thermodynamic cycles connecting receptor conformations. These thermodynamic cycles (Figures 1 and 5) resemble the pharmacological representation of GPCR activation, which are the basis for the estimation of the corresponding equilibrium constants (10). In our case, we compare the activation pathway between chemically related species, these being either pairs of ligands or single-point mutants *vs* the wt receptor. Taking additional advantage of the increased structural information of GPCRs in different conformations, as is the case of the A_2A_AR, one can solve the vertical legs of the designed cycles via automated FEP protocols tailored for ligand (25) or residue (26) perturbations, respectively. Following the scheme depicted in Figure 1B, we demonstrate the validity of this approach with the calculation of structure-efficacy relationships for three series of A_2A_AR ligands, resulting in excellent discrimination of full and partial agonists from neutral antagonists (Figures 3 and 4). The agonistic profile is modeled as the capacity of a molecule to achieve the desired conformational selectivity for the active configuration, opening the door to the structure-based computational design of compounds with tailored pharmacology, with additional potential to provide insights in pharmacogenomics of drugs (37).

Using the analogous approach depicted in Figure 1C, we show how to estimate the effect of protein mutations on the conformational equilibrium of the receptor, which can be translated to variations on its basal activity. The approach is applied to characterize a constitutively inactive mutation (CIM, Thr88^3.36^A) as well as a neutral mutation (Ser277^7.43^A), finding the same discrimination as the experimental data available in both cases (Figure 5B). Moreover, taking advantage of the additive property of the thermodynamic cycles, one can model ligand efficacy as the estimation of the ligand affinity for the active (R*) conformation weighted by the accessibility of the ligand to R*. The combined cycle, shown in Figure 5A, allowed a successful discrimination of the different effects exerted by each of these two mutations on the efficacy of a full and a partial agonist. The relevance of this result goes beyond the successful correlation of the computed ranking of the mutational shifts in ligand efficacy with the experimental values, and additionally provides a structural and mechanistic framework to interpret these results. Thus, it was found that the CIM T88^3.36^A, by displacing the equilibrium towards the inactive conformation, can neutralize the predicted increase in affinity for the partial agonist LUF5834 or, conversely, display a synergistic effect on the predicted decrease in the R* affinity for the full agonist CGS21680, explaining the dramatic decrease on its efficacy for the A_2A_AR (Figure 5B). The mutation S277^7.43^A was found to be neutral with regards to the receptor conformational equilibrium, in agreement with its negligible effect on the basal activity of the receptor. Consequently, the correctly predicted net effect in increasing or decreasing the ligand efficacy of the partial and full agonist, respectively, is directly correlated to the predicted effects of the mutation on the affinity for R* in each case (Figure 5B).

The presented approach provides a generally applicable framework, as long as the underlying pharmacological problem can be based on clear structural endpoints (in the present study, an active and inactive structure of the receptor). The rapidly expanding arsenal of GPCR structures (38), with various states available for a number of receptors (16,39), will widen the applicability of our workflow to cover a large extent of the GPCR-ome. These structures not only include G protein ternary complexes, but lately also e.g. β-arrestin bound structures (40), or nanobody derived structures of intracellular binding partners (41). As such, the method is not limited to the prediction of agonist profiles of compounds, but could potentially be extended to the design of biased ligands(42). Moreover, there is now strong evidence from HDX (43) or NMR (44) experiments of intermediate conformational states of GPCRs, including recent proposals that the partial agonists of the A_2A_AR could preferentially bind one of these intermediate states with compromised G-protein coupling (44,45). One could expect that the precise structures of those states could be characterized in the near future, and consequentially use them for building more precise thermodynamic cycles representing revised pharmacological models of receptor activation(44,46). Alternatively, the derivation of activation states from inactive receptors via MD approaches can provide structural insights into receptor activation mechanisms (47), and identify intermediate states along the receptor activation path. In addition, the experimental structures available provide excellent starting points for the generation of reliable homology models (48), which can be also useful starting structures for FEP simulations, as we recently showed in the case of the orphan GPR139 (49), the neuropeptide Y receptor family (26,50) or the A_2B_AR (51).

To the best of our knowledge, this is the first time that the activation of a GPCR is modeled with the use of thermodynamic cycles coupled to FEP simulations. The generated framework is easily extensible to other GPCRs, offering a computational approach to design ligands with tailored pharmacological properties or to predict the effect of point mutations on the receptor conformational equilibrium. The last question is gaining interest as we see increasing examples of GPCR point mutations related to disease, as is the case of the A_2B_AR in cancer (52), by the mechanism of shifting the basal activity to either constitutively inactive or constitutively active mutations. An additional advantage of our end-point approach, besides the computational efficiency, is its modularity, allowing the combination of thermodynamic cycles to predict e.g. shifts in ligand efficacy induced by point mutations. Moreover, the protocol could potentially be extended to, e.g., DFG in and out kinases (53), open and closed state of ion channels(54) or single solute carrier (SLC) transporters (55), provided that the chemical space of the studied cases sufficiently overlap (i.e. point mutations or related chemotypes).

## Methods

### Structure preparation, membrane insertion and equilibration

The high-resolution crystal structure of the A_2_AR with antagonist ZM241385 [4EIY (12)] was used as receptor starting structure for the inactive state, whereas 2YDV (31) and 5G53 (14) were used for the active state. Thereafter, the engineered BRIL fusion protein was removed and missing loops (C-terminal fragment of EL2 and most of EL3) were modelled, and protonation states of residues assigned, as described elsewhere (56). The active structure was thereafter aligned to the inactive structure of the receptor. The 3D coordinates for the ligands were generated with LigPrep and subsequently docked to the prepared A_2A_AR structure using Glide (57), and the antagonists structures by flexible ligand alignment to their reference agonist compound. Apo structures were generated by removing the ligand, but keeping crystallographic waters, and subsequently embedded in a solvated membrane environment using PyMemDyn (58). Shortly, this protocol embeds a structure in a pre-equilibrated POPC (1-palmitoyl-2-oleoyl phosphatidylcholine) membrane model such that the TM bundle is parallel to the vertical axis of the membrane. The system is then soaked with bulk water and inserted into a hexagonal prism-shaped box that is energy-minimized and carefully equilibrated during 5 ns, following the PyMemDyn protocol described elsewhere (58). The standard OPLS all-atom (OPLS-AA) force field is used for all residues (59), and parameters for membrane lipids were taken from the Berger united-atom model (60). The corresponding equilibrated holo structures were generated by restoring the docked ligands and removing overlapping water molecules.

### FEP simulations

The receptor-ligand structures in the equilibrated membrane were subsequently transferred to the MD package Q, in order to perform FEP calculations under spherical boundary conditions (61). A 50 Å diameter sphere was centered on the center of geometry of ZM241385 (or equivalent point in the remaining structures), where solvent atoms are subject to polarization and radial restrains using the surface constrained all-atom solvent (SCAAS) model to mimic the properties of bulk water at the sphere surface (62). Atoms lying outside the simulation sphere are tightly constrained (200 kcal/mol/Å^2^ force constant) and excluded from the calculation of non-bonded interactions. Within the simulation sphere, long range electrostatics interactions beyond a 10 Å cut off were treated with the local reaction field method (63), except for the atoms undergoing the FEP transformation, where no cutoff was applied. Solvent bond and angles were constrained using the SHAKE algorithm (64). All ionizable residues outside the sphere and those within the boundary were considered in their neutral form as described elsewhere (25). Residue parameters were translated from the latest version of the OPLSAA/M force field (59), whereas ligand OPLS2005 parameters were retrieved from Schrodinger’s ffld_server (65), and translated to Q following the QligFEP protocol (25). The simulation sphere was heated from 0.1 to 298 K during a first equilibration period of 0.61 nanoseconds, where the initial restraint of 25 kcal/mol/Å^2^ imposed on all heavy atoms was slowly released. Thereafter the system was subject to unrestrained MD simulations, starting with a 0.25 nanosecond unbiased equilibration period which is followed by the FEP sampling, following different protocols for sidechain and ligand perturbations as detailed below. In both cases, atom transformations occur between initial and ending states, evenly divided into a number of λ windows that depend on the FEP protocol adopted (see below). This sampling is replicated in 10 independent MD simulations with different initial velocities, each of them consisting of 10 ps sampling per λ window using a 1 fs time step in all cases.

The generalized FEP protocol for amino acid mutations, QresFEP (26,66) was used to estimate the effects of single point mutations on ligand binding. Briefly, QresFEP is a single-topology FEP protocol that divides the sidechain perturbation to alanine into separate stages, where atom annihilations occur gradually for each charge group (as defined on the OPLS force field), starting from the most topologically distant from the C_β_ atom, in four consecutive stages (66): 1) the partial charges are initially removed, 2) van der Waals potentials are transformed into smoother soft-core potentials, 3) annihilation of the corresponding group of atoms., 4) restoring the partial charges of the final species. The number of perturbation stages needed for the full annihilation depends on the nature of the sidechain involved, in this case ranging from four (Ser) to five (Thr), where each of the subsequent stages is evenly divided into 20 λ windows. To fulfill a thermodynamic cycle, the same sidechain annihilation is simulated in the apo structure of the protein, so that the energetic difference between these two processes equals the binding affinity shift due to the mutation. It follows that the sampling time for the sidechain perturbation here considered was 8-10 ns.

All ligand perturbations were performed with our dual-topology QligFEP protocol (25), where the full transformation of one ligand into another is performed along a linear λ sampling consisting of 50 windows. The reference state to create the thermodynamic cycle is in this case the MD simulation of the ligands’ transformation in an equivalent sphere of water, and the in this case difference between the protein-bound and reference ligand transformations equals the difference in binding affinity between the two ligands.

In both residue and ligand transformations, relative binding free energies were estimated by solving the thermodynamic cycle utilizing the Bennet acceptance ratio method (BAR) (67) as

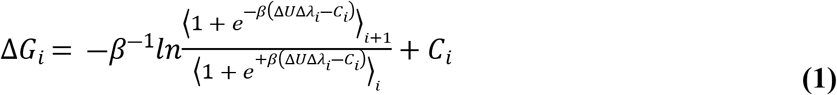

where the constants *C_i_* are optimized iteratively so that the two ensemble averages become equal, yielding *ΔG_i_* = *C_i_*. Average values ± SEM are reported from the 10 independent MD simulations in each relevant state.

## Acknowledgements

The computations were performed on resources provided by the Swedish National Infrastructure for Computing (SNIC). Additional support from the Swedish strategic research programme eSSENCE is acknowledged. The authors participate in the European COST Action CA18133 (ERNEST).

